# High-throughput microplate luminometry of *Pyrocystis lunula* bioluminescence for metal toxicity assessment

**DOI:** 10.1101/2025.10.22.682404

**Authors:** L.S. Perin, F. M. Saldanha-Corrêa, M. Menashe, L. Rahmani, E. S. Braga, A. G. Oliveira

## Abstract

Dinoflagellates are major contributors to marine phytoplankton, and many species emit blue bioluminescence from specialized organelles called scintillons in response to mechanical stimulation and transient intracellular acidification. Because pollutants can disrupt this pathway, changes in light output provide a sensitive proxy for environmental stress. We utilized the bioluminescence of dinoflagellate *Pyrocystis lunula* to develop a rapid, high-throughput toxicity assay. Cultures aged 30-40 days were exposed to a range of metal concentrations for 24 hours, and stimulus-evoked emission was quantified in 96-well plates using a microplate luminometer. All metals tested produced dose-dependent inhibition of bioluminescence, with sensitivity varying by contaminant. Non-essential metals showed the strongest inhibitory effects: Cd^2+^ (half-maximal inhibitory concentration, IC_50_, 24 h = 0.014 mg L^−1^) and Pb^2+^ (IC_50_, 24 h = 0.016 mg L^−1^). Essential metals, Cu^2+^ and Zn^2+^, also reduced emission but at higher concentrations; differences were significant by one-way analysis of variance with Tukey’s post hoc test. This assay is rapid, low-cost, and scalable, offering a practical tool for monitoring trace-metal contamination and supporting ecotoxicological assessments.

## INTRODUCTION

Bioluminescence, the emission of visible light through a chemical reaction by living organisms, is widespread among marine animals and microorganisms (Herring 1987; Shimomura 2006; Moline et al. 2007; Haddock et al. 2010; Lau and Oakley 2021; Claes et al. 2024). Chemically, a bioluminescent reaction is an exergonic oxidation of a substrate, generically called a luciferin, by molecular oxygen and is catalyzed by enzymes known as luciferases, producing visible photons (Wilson and Hastings 1998; Shimomura 2006; Haddock et al. 2010; Lau and Oakley 2021; Schramm and Weiß 2024). Elucidation of these mechanisms has advanced enzymology and evolutionary biology and has enabled broad biotechnological applications (Ogawa et al. 1995; Gerdes and Kaether 1996; Girotti et al. 2008; Syed and Anderson 2021). In environmental science, bioluminescence is used primarily in ecotoxicology to assess the effects of natural and synthetic substances on organisms, populations, and communities (Truhaut 1977; Zagatto and Bertoletti 2006). Common assays expose naturally luminous or engineered bioluminescent bacteria to environmental samples, diluted for quantification or undiluted for screening, and compare their light output with controls, as in the Microtox^®^ test (Zagatto and Bertoletti 2006; Girotti et al. 2008). *Aliivibrio fischeri* (formerly *Vibrio fischeri*) is the primary test organism for these assays, which are widely applied to effluents, waters, and sediment extracts. The assays are rapid, reproducible, and cost-effective, making them suitable for both preliminary screening and in-depth evaluations, and they are standardized by regulatory agencies (EPA 2013; ABNT 2021a, b, c).

In addition to bacterial systems, bioluminescence-based toxicity assays have been applied, although less extensively, to luminous dinoflagellates (Hannan et al. 1986; Lapota et al. 1993, 1995; Okamoto et al. 1999; Heimann et al. 2002; Craig et al. 2003; Rosen et al. 2008; Stauber et al. 2008; Hildenbrand et al. 2015). Dinoflagellates are the principal eukaryotic protists capable of producing light (Haddock et al. 2010; Hastings 2013). Dinoflagellate bioluminescence occurs within specialized organelles called scintillons through a luciferin-luciferase reaction, producing blue flashes (~480 nm) that last a few seconds (Shimomura 2006; Haddock et al. 2010; Valiadi and Iglesias-Rodriguez 2013; Fajardo et al. 2019; Fajardo et al. 2020). Dinoflagellates are unicellular organisms that may be photosynthetic, heterotrophic, or mixotrophic; they are cosmopolitan and ecologically important members of marine phytoplankton, often engage in symbiotic or parasitic relationships, and include taxa capable of producing toxins (Taylor 1987). As primary producers, dinoflagellates can provide early indicators of impacts on marine biota and ecosystem services. Using a eukaryotic model may improve the predictive value for potential human toxic effects compared with bacterial tests (Hannan et al. 1986; Lapota et al. 1993, 1995; Okamoto et al. 1999; Heimann et al. 2002; Craig et al. 2003; Rosen et al. 2008; Stauber et al. 2008; Hildenbrand et al. 2015; Perin et al. 2022).

Although the mechanisms underlying dinoflagellate bioluminescence are not yet fully resolved, applications in ecotoxicological assays already show a clear dose-dependent light response, with performance that compares well to conventional tests using fish, invertebrates, other algae, and bacteria. These assays also stand out for their low cost, ease of implementation, and rapid turnaround, which make them attractive for routine monitoring programs (Hannan et al. 1986; Lapota et al. 1993, 1995; Okamoto et al. 1999; Heimann et al. 2002; Craig et al. 2003; Rosen et al. 2008; Stauber et al. 2008; Hildenbrand et al. 2015). Despite these strengths, and the growing need for scalable monitoring tools amid intensifying anthropogenic pressures on marine ecosystems, the ecotoxicological use of dinoflagellate bioluminescence remains underutilized. This limited application is particularly significant given the challenge of monitoring trace metals. Some metals, such as copper and zinc, are essential at low concentrations because they serve critical physiological roles, yet their environmental persistence, bioaccumulation, and biomagnification can drive exposures into harmful ranges. Other metals, including cadmium and lead, lack biological function and are toxic at concentrations only modestly above natural background levels. As a result, metals represent a priority class of contaminants for which sensitive, rapid, and cost-effective bioassays are urgently needed, and dinoflagellate bioluminescence provides a mechanistically relevant and operationally practical platform to meet this need (Raven et al. 1999; Omar 2002; Travieso et al. 2002; Islam and Tanaka 2004; Sharma and Agrawal 2005; Visbeck 2018; Le et al. 2024; Allan 1997; Stern 2010; Schoofs et al. 2024).

The limited use of bioluminescent dinoflagellates in environmental testing may stem from the low analytical capacity of traditional methodologies, typically restricted to just 1–6 samples per run. Here, we propose a new ecotoxicological assay that incorporates updated measurement technology using a microplate luminometer, which optimizes data acquisition and enables the simultaneous analysis of multiple samples. This approach improves applicability, provides an additional tool for toxicity testing, and aligns with ecotoxicological guidelines, which emphasizes that a reliable assessment of chemical toxicity on the biota requires at least three bioassays with organisms from different trophic levels (Rand 1995; Zagatto and Bertoletti 2006; Magalhães and Ferrão-Filho 2008). By introducing an updated methodology that enhances the use of dinoflagellates as bioindicators, this study contributes both to addressing regulatory requirements and to providing an additional approach to ecotoxicological testing.

## MATERIALS & METHODS

### Cultures of *Pyrocystis lunula*

The dinoflagellate *Pyrocystis lunula* (Schütt, 1896) was obtained from the Aidar & Kutner Microorganism Bank (BMAK; strain BMAK 264) located at Oceanographic Institute, University of São Paulo. Cultures were grown in 300 mL Erlenmeyer flasks in an incubator (Caltech, EI-08F1/F2-F) at 21 ± 1 °C under a 12:12 h light:dark cycle (irradiance 110 µmol photons m^−2^ s^−1^). Cell density and culture health were assessed weekly by light microscopy and hemocytometer counts until cultures were 25–30 days old (for inoculation) or 30–40 days old (for exposure assays). Guillard’s f/2 medium (Guillard and Ryther 1962; Guillard 1975) was prepared with reagent-grade chemicals (Sigma-Aldrich) and adjusted to a salinity of 35. Each culture was initiated at a volume of 250 mL with an initial cell density of 1,000 cells mL^−1^. All inoculation and exposure procedures were performed in a laminar-flow cabinet (Pachane, PA420) in low light conditions.

### Metal Exposure and Bioluminescence Assay in *Pyrocystis lunula*

#### Experimental setup

Cultures aged 30-40 days were used. For each exposure, tubes contained 3.0 mL at 3,000 cells mL^−1^ (9,000 cells total), which corresponds to 300 cells per 100 µL (the assay volume per well). Cell counts were performed on the dosing day to verify the density of the stock cultures and to calculate the aliquot volume required to reach the target density in the exposure. Culture addition was limited to 2.5 mL. When necessary, volumes were adjusted with Guillard’s f/2 medium standardized at a salinity of 35 to maintain consistency across all conditions.

#### Exposure

Analytical-grade salts from Sigma-Aldrich were used: CuCl_2_, CdCl_2_·2.5H_2_O, ZnSO_4_·7H_2_O, and Pb(NO_3_)_2_. Each metal stock solution (500 mL) was prepared at 0.3 mM using saline solution (NaCl) at a salinity of 35. Test concentrations were 0.5, 1.0, 3.0, 5.0, 7.0, and 10.0 µM, plus a no-metal control. The stock solution was diluted in Guillard’s f/2 medium to reach the target final exposure concentrations, and up to 0.5 mL of this mixture was added to 2.5 mL of culture. Dosing was performed 4 h into the dark phase. Each condition was prepared in biological triplicate (n=3). Tubes were incubated for 24 h under the same temperature and photoperiod as the maintenance cultures.

#### Bioluminescence measurement

After 24 h exposure, four aliquots of 100 µL from each tube were transferred to separate wells of a black 96-well plate. Bioluminescence was elicited by injecting 100 µL of fresh Guillard’s f/2 medium (salinity 35) into each well via a Tecan Infinite^®^ 200 luminometer injector, followed by a 10 s integrated read. Plates were dark-adapted in the instrument for 10 min before the first measurement, and wells were read individually.

#### Data Analysis and Statistical Treatment

Data generated by the Magellan^®^ software were evaluated for normality using the Shapiro–Wilk test. Data that passed the normality test were analyzed using one-way ANOVA followed by Tukey’s post hoc test, using GraphPad Prism (version 10.3.0). Mean bioluminescence values for each concentration were calculated using Excel^®^, and outliers were removed using the interquartile range method (IQR): values above Q3 + 1.5 × IQR or below Q1 – 1.5 × IQR were discarded prior to averaging (Fávero and Belfiore 2017; Vinutha et al. 2018). Bioluminescence inhibition was expressed as a percentage, with the control (no contaminant) defined as 100%. To determine the toxicity of metals and facilitate their quantitative comparison, the IC_50_ values were estimated using a four-parameter logistic model (log[inhibitor] vs. normalized response) in GraphPad Prism. The experimental design for the assay employing a bioluminescent primary producer bioindicator and microplate luminometers is depicted in Figure 1.

**Fig. 1.**
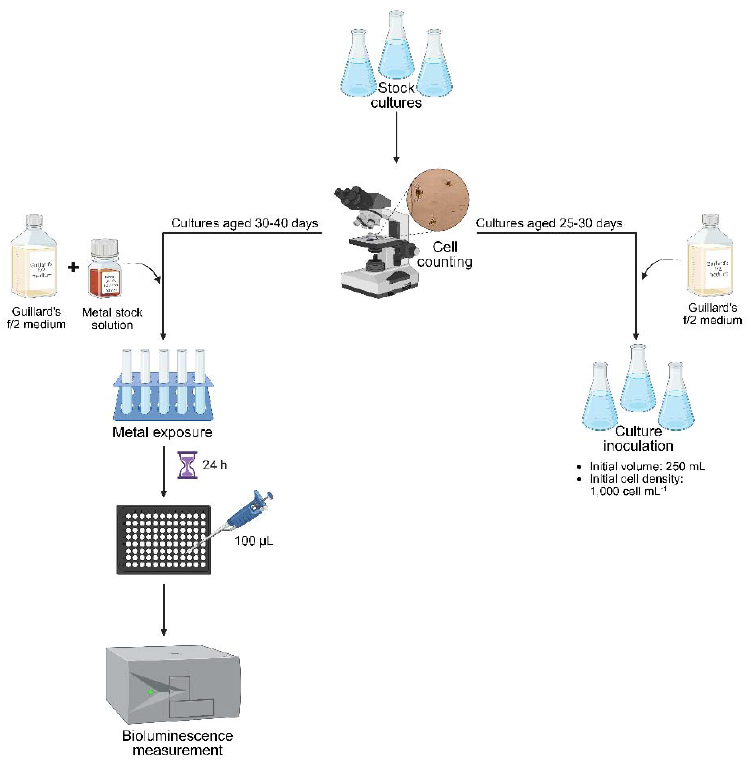
Experimental workflow for dinoflagellate bioluminescence assays under metal exposure. Stock cultures were maintained in Guillard’s f/2 medium and monitored by cell counting. Cultures aged 25–30 days were used for inoculation (initial volume 250 mL; 1,000 cells mL^−1^), while cultures aged 30–40 days were exposed to metal stock solutions. After 24 h exposure, 100 µL of each treatment was transferred to a microplate and bioluminescence was quantified using a luminometer. Created with BioRender.com/wacyrug

## RESULTS

### Optimization of experimental conditions

The use of bioindicators has become essential for environmental monitoring given the complexity of interactions between chemical agents and biological effects, as well as the limitations of purely physico-chemical methods (e.g., chromatography and spectrometry) in detecting such effects. Here, we describe an alternative toxicity-assessment method using microplate luminometry for measuring light variation from a bioluminescent marine primary producer, expanding the set of practical bioindicators. To stimulate dinoflagellate bioluminescence in the luminometer, injection parameters had to be experimentally determined. Volumes of 5, 20, and 100 µL were tested (considering well capacity). Light-emission data met normality by the Shapiro–Wilk test. One-way ANOVA followed by Tukey’s test showed significantly higher stimulation at 100 µL (5 µL vs. 100 µL, p = 0.0091; 20 µL vs. 100 µL, p = 0.0304; 5 µL vs. 20 µL, p = 0.5572; Fig. 2a). Regarding the injected liquid, we compared 100 µL of culture medium versus 100 µL sodium citrate, given that dinoflagellate bioluminescence depends on scintillon acidification and prior studies have used acids to trigger light emission (Okamoto et al. 1999; Shimomura 2006). Emission data again met normality (Shapiro–Wilk), and one-way ANOVA detected no significant difference between liquids (p = 0.9929; Fig. 2b).

**Fig. 2.**
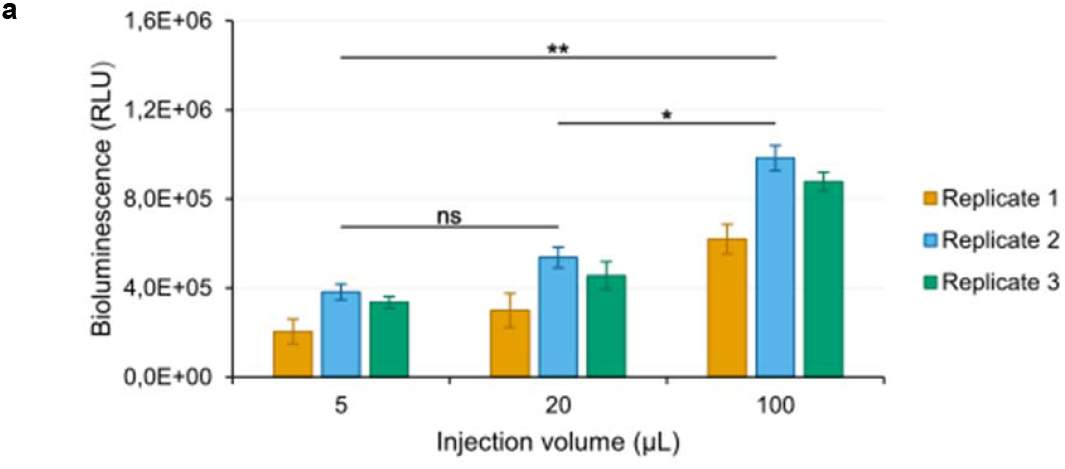
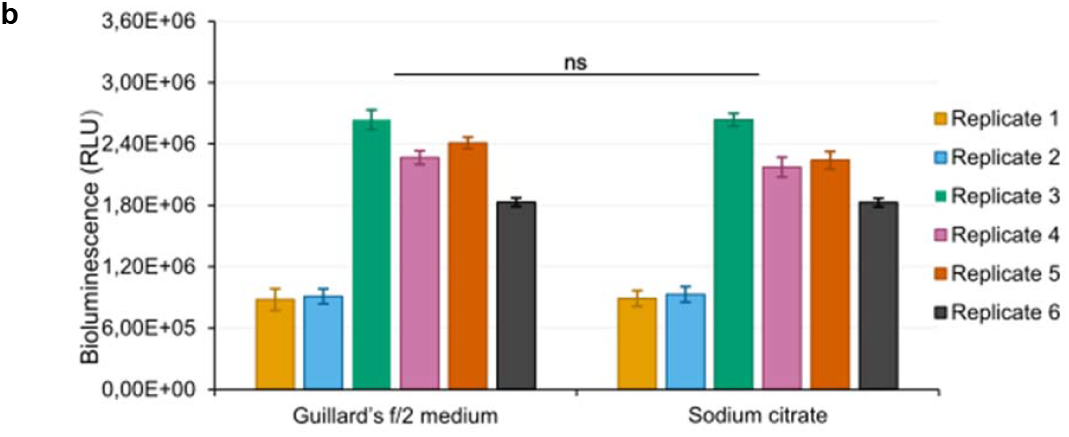
Bioluminescence of *Pyrocystis lunula* in response to **a**) different injection volumes used to generate the mechanical stimulus; **b**) different injection solutions used to generate the mechanical stimulus. Asterisk indicates a significant difference by Tukey’s; (*) at p = 0.0304; (**) at p ≤ 0.0091; (ns) indicates no significant difference. Error bars represent the standard deviation of the dataset.

### Effect of metals on bioluminescence

Box plots in Figure 3 summarize *P. lunula* bioluminescence after 24 h exposures to four trace metals (Zn^2+^, Cu^2+^, Cd^2+^, Pb^2+^). Across all metals, we observed a concentration-dependent inhibition of light emission. At the lowest doses (0.5 and 1.0 µM) and in the controls, variability was greater, evidenced by elongated boxes and more outliers. Normality of the data was confirmed by Shapiro–Wilk tests. Consequently, we applied one-way ANOVA followed by Tukey’s post hoc test to evaluate the effects of metal concentration on bioluminescence inhibition. ANOVA indicated a significant concentration effect for Zn^2+^, Cu^2+^, Cd^2+^, and Pb^2+^ (p < 0.001). For Cu^2+^ and Zn^2+^, post hoc tests showed that the three highest concentrations differed significantly from their run-specific controls, whereas the three lowest concentrations (0.5, 1.0, and 3.0 µM) did not (p > 0.05). In contrast, for Cd^2+^ and Pb^2+^, all tested concentrations differed significantly from their respective controls (p < 0.05; Fig. 3). Individual boxplots for each test replicate are provided in the supplementary materials.

**Fig. 3.**
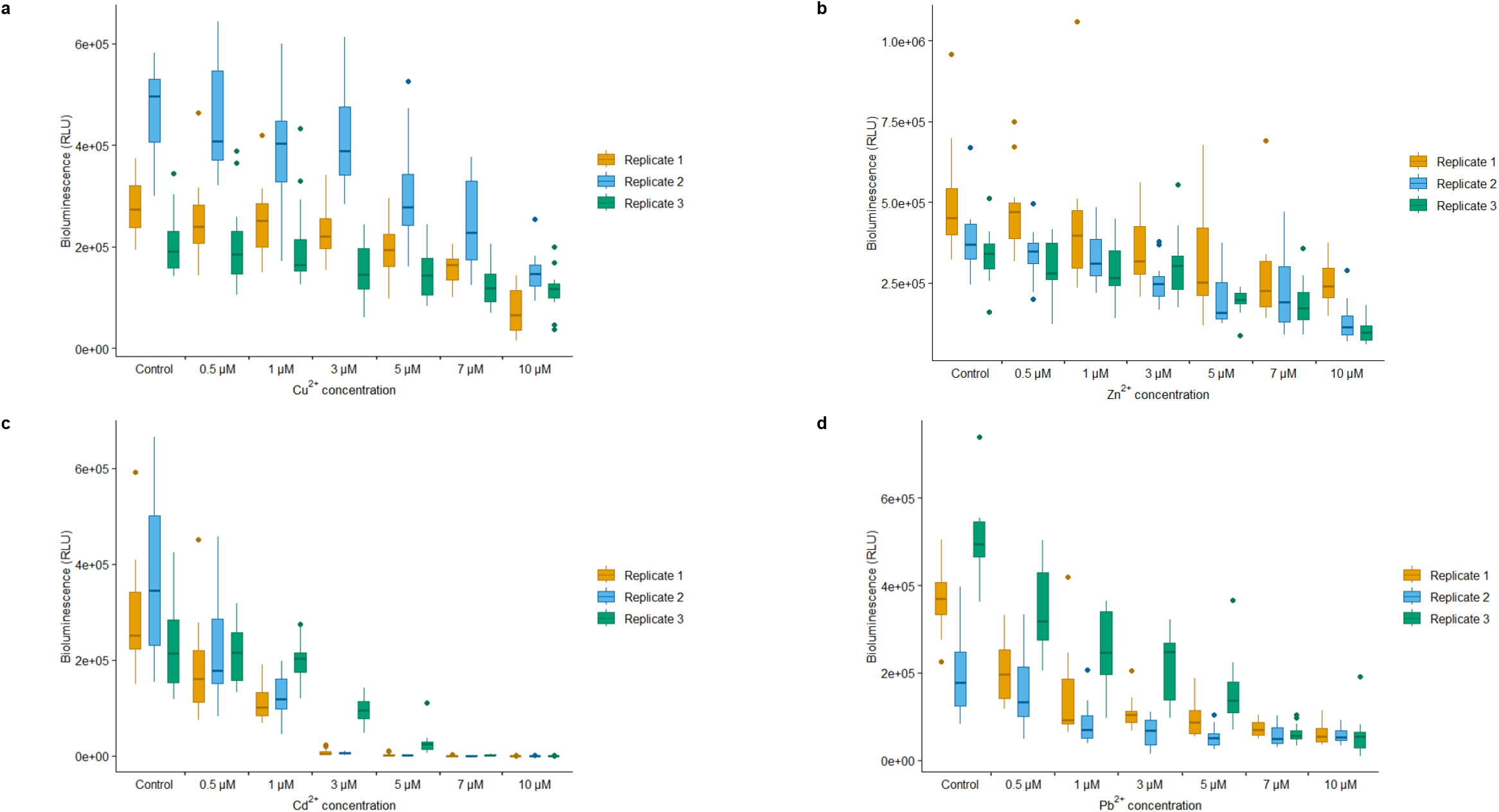
Light emission of *P. lunula* at different concentrations of **a**) Cu^2+^; **b**) Zn^2+^; **c**) Cd^2+^; **d**) Pb^2+^.

Percent inhibition of bioluminescence was calculated relative to the concurrent control in each ecotoxicological assay (Fig. 4). A clear dose-dependent effect was observed: *P. lunula* luminescence decreased progressively with increasing metal concentration. Nonessential metals (Cd^2+^, Pb^2+^) produced stronger inhibition, with >50% reduction already at 1.0 µM and near-complete suppression by Cd^2+^ at 5.0-10.0 µM. By contrast, the essential metals Zn^2+^ and Cu^2+^ caused marked inhibition only at higher concentrations; at 10.0 µM, bioluminescence was reduced by >50%.

**Fig. 4.**
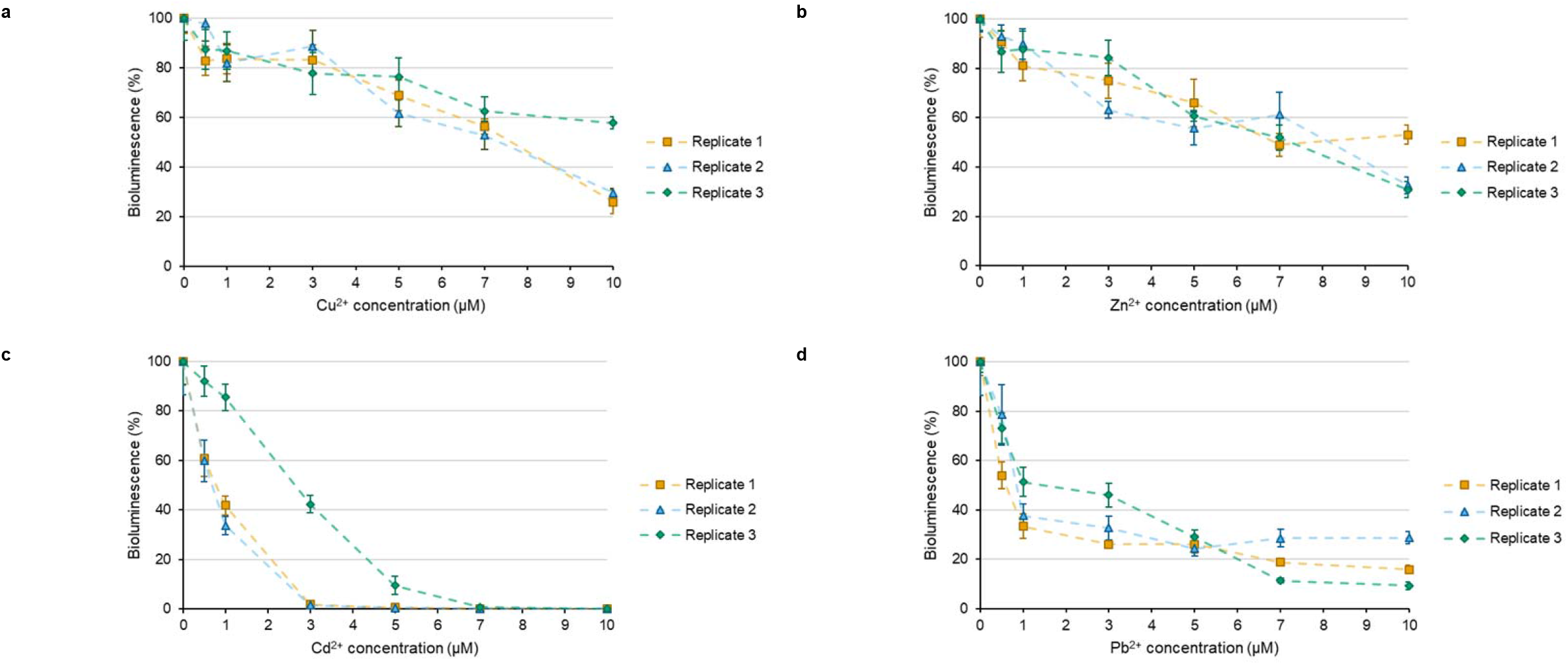
Effects of increasing trace metal concentrations on the bioluminescence of the dinoflagellate *Pyrocystis lunula*. **a**) assays with Cu^2+^; **b**) assays with Zn^2+^; **c**) assays with Cd^2+^; **d**) assays with Pb^2+^. Error bars represent the standard error of the dataset.

The IC_50_ parameter is widely used to quantify the potency or toxicity of substances and mixtures in inhibiting biological or biochemical processes (Zagatto and Bertoletti 2006; Sebaugh 2011). In our *P. lunula* assays, 24 h IC_50_ values for Cd^2+^ and Pb^2+^ were similarly low, indicating high sensitivity to these contaminants: Cd^2+^ yielded an IC_50_-24 h of 0.014 mg L^−1^ (95% CI: 0.008–0.024 mg L^−1^), and Pb^2+^ an IC_50_-24 h of 0.016 mg L^−1^ (95% CI: 0.010–0.023 mg L^−1^). For both, coefficients of determination (R^2^) exceeded 0.9, indicating good agreement between the data and the selected dose-response model. By contrast, IC_50_-24 h values could not be estimated for Cu^2+^ and Zn^2+^ within the tested range (0.003–0.064 mg L^−1^), because inhibition did not exceed 50%, resulting in a poor model fit. As recommended for reliable IC_50_ estimation, at least two concentrations producing >50% inhibition are needed (Sebaugh 2011). Given that concentrations ≥10.0 µM of Cu^2+^ and Zn^2+^ produced >50% inhibition in our experiments, future tests should include concentrations above 10.0 µM for these essential metals.

## DISCUSSION

### Methodological considerations of the proposed protocol

Pollution events are progressively affecting the marine environment, underscoring the need for diverse monitoring tools to adequately assess their impacts. As an environmental indicator, a basal-trophic bioluminescent organism was chosen for its ecological relevance, sensitivity to environmental substances, and easily detectable physiological response. Photosynthetic dinoflagellates such as *P. lunula* play a key role in oceanic primary production and thus in the flow of matter and energy; their development depends on environmental factors including light, temperature, salinity, and nutrient concentrations (Taylor 1987). Accordingly, *P. lunula* can provide early responses to potential stressors on marine biota and ocean ecosystem services. As a bioluminescent species, it also offers a readily measurable sublethal physiological endpoint that has been successfully used in previous bioassays (Hannan et al. 1986; Lapota et al. 1993; Heimann et al. 2002; Craig et al. 2003; Rosen et al. 2008; Stauber et al. 2008; Hildenbrand et al. 2015), allowing comparison with existing results. As a sublethal indicator, changes in luminescence provide an early warning of adverse effects in the organism, enabling the detection of impacts before morphologically visible or lethal damage occurs. This approach supports the establishment of more protective environmental quality criteria, helping to prevent irreversible effects and maintain the long-term viability of populations (Perkins, 1979; Zagatto and Bertoletti, 2006). Additional advantages include wide geographic distribution (enabling deployment across regions), small size, and ease of culture, which facilitate laboratory use (Zagatto and Bertoletti, 2006). Cultures were used for contaminant exposure at 30–40 days of age to ensure use in the stationary phase and to reach the minimum cell density required for the experiments. Because effects were measured 24 h after exposure and bioluminescence readings were set to 300 *P. lunula* cells per 100 µL measurement aliquot, it was important that cell concentrations remained stable throughout testing. The choice of 300 cells per 100 µL was based on Lapota et al. (1993), who evaluated light output as a function of cell number and observed that increasing cell concentration (and thus light emission) reduced the coefficient of variation (CV) among triplicates; at 300 cells the CV was ~10%, indicating low variability. To achieve this density, 9,000 cells were inoculated in 3 mL, requiring stock cultures to reach ≥3,600 cells mL^−1^ so that dinoflagellate aliquots could be added up to 2.5 mL of exposure solution. Exposures began 4 h after the onset of the dark phase, when light responses are maximized and consistent (Stauber et al. 2008); a pattern also confirmed in our experiments. To improve throughput and reproducibility, we adopted a microplate luminometer equipped with an injection system. Earlier studies employing dinoflagellates typically measured 1–6 cuvettes at a time, necessitating multiple reading rounds or reduced sample numbers (Hannan et al. 1986; Lapota et al. 1993, 1995; Okamoto et al. 1999; Heimann et al. 2002; Craig et al. 2003; Rosen et al. 2008; Stauber et al. 2008; Hildenbrand et al. 2015). Plate-based readings with per-well injection enable stimulation and detection in each well with minimal manual intervention, allowing simultaneous testing of multiple samples and triplicates and thereby increasing efficiency.

We selected injector-based stimulation rather than plate shaking to optimize and standardize the mechanical stimulus. Shaking affects the entire plate simultaneously and cannot deliver controlled, per-well stimulation; wells read later may experience repeated stimuli, potentially diminishing bioluminescence due to accumulated stress and increasing variability. As mentioned earlier, the selected injection volume was 100 µL, which corresponds to the maximum volume permitted by the equipment and matches the exposure aliquot added to each well. This volume was chosen because it produced the strongest bioluminescent response. Since no significant differences were observed between the tested liquids, the response appears to be triggered mostly by mechanical impact, not by changes in pH, and Guillard’s f/2 medium was therefore selected.

### Ecotoxicological assays

This study shows that bioluminescence inhibition by exposure to the selected metals can be quantified with good reproducibility. All metals produced dose-dependent responses; however, response profiles differed between essential and nonessential metals and across the tested concentration ranges. At the lowest exposures (0.5 and 1.0 µM) and in the controls, data variability was greater, likely reflecting experimental noise and intrinsic biological heterogeneity. Intraspecific variation in light production is expected because dinoflagellate bioluminescence is influenced by physiological and environmental factors (Shimomura 2006; Timsit et al. 2021; Claes et al. 2024), as evident in the variability among control groups. The pronounced variability at the lowest metal concentrations suggests that toxicity at these levels is insufficient to induce a uniform inhibitory response. This is consistent with essential metals, such as Cu^2+^ and Zn^2+^, which play key biological roles and therefore can elicit more heterogeneous responses, as reflected by the higher number of outliers for these contaminants.

Prior work with bioluminescent dinoflagellates has similarly documented deleterious effects of these metals on light emission (Lapota et al. 1993, 1995; Okamoto et al. 1999; Heimann et al. 2002; Craig et al. 2003; Rosen et al. 2008; Stauber et al. 2008). The precise mechanism by which metals suppress bioluminescence remains unresolved. In *Lingulodinium polyedrum*, Okamoto et al. (1999) observed an immediate increase in spontaneous flashes after the addition of Hg^2+^, Cd^2+^, Pb^2+^, or Cu^2+^ and proposed that metals raise the frequency of membrane action potentials, triggering scintillon acidification and, consequently, light emission.

Toxic metal ions can enter cells by forming complexes with membrane lipids/proteins, by co-opting multivalent ion transporters that normally import essential ions, or via ionic mimicry (e.g., Cd^2+^/Pb^2+^ mimicking Ca^2+^). These processes can depolarize the membrane and acidify the cytoplasm (Green et al. 1980; Okamoto et al. 1999; Pinto et al. 2003; Singh et al. 2016). Building on this rationale, Heimann et al. (2002) examined bioluminescence recovery in *P. lunula* exposed to Cu^2+^, Cd^2+^, Pb^2+^, Ni^2+^, sodium dodecyl sulfate (SDS), phenol, and phenanthrene. They proposed that metals stimulate light emission during the first ~4 h via scintillon acidification, partially depleting the bioluminescent system and producing a dose-dependent “recovery” pattern when emission is measured 4 h after exposure.

IC_50_ values were used to compare the results obtained with the proposed toxicity assay protocol to those reported in previous studies employing dinoflagellate bioluminescence in environmental testing, as well as to other bioindicators commonly applied in ecotoxicology. This comparison situates the sensitivity observed in *P. lunula* with findings from earlier works (Lapota et al. 1993, 1995; Heimann et al. 2002; Rosen et al. 2008; Stauber et al. 2008) and highlights similarities and differences relative to traditional model organisms, such as *A. fischeri, Daphnia magna* and *Hyalella azteca*. Figure 5 summarizes these comparisons, presenting the IC_50_ values obtained in this study (diamonds) alongside literature data of Cu^2+^, Zn^2+^, Cd^2+^, and Pb^2+^.

**Fig. 5.**
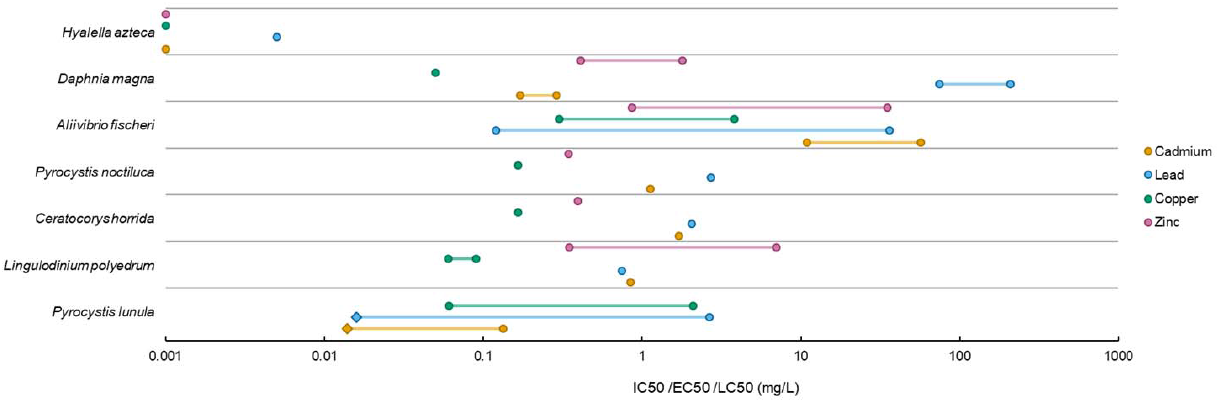
Acute effects of trace metals Cd^2+^, Pb^2+^, Cu^2+^, and Zn^2+^ in different ecotoxicological assays. IC_50_ values obtained in this study are indicated by diamonds.

For dinoflagellate assays, the IC_50_ values obtained using our methodology were generally lower than previously reported. For Cd^2+^, *P. lunula* showed sensitivity closer to results reported by Heimann et al. (2002; IC_50_-24 h = 0.133 mg L^−1^) than to Rosen et al. (2008), who reported that *L. polyedrum* (IC_50_-24 h = 0.843 mg L^−1^) was more sensitive than *Ceratocorys horrida* (IC_50_-24 h = 1.710 mg L^−1^), and *Pyrocystis noctiluca* (IC_50_-24 h = 1.130 mg L^−1^), indicating interspecific differences in sensitivity (Fig. 5).

For Pb^2+^, *P. lunula* was likewise more sensitive than the species examined by Rosen et al. (2008), including *L. polyedrum* (IC_50_-24 h = 0.747 mg L^−1^), *C. horrida* (2.060 mg L^−1^), and *P. noctiluca* (2.710 mg L^−1^). The IC_50_ measured in this study was markedly lower than that reported for *P. lunula* by Heimann et al. (2002; IC_50_-24 h = 2.645 mg L^−1^), consistent with mechanisms that require longer exposure to elicit effects at lower concentrations (Trevors et al. 1986; Nowicka 2022; Le et al. 2024).

Considering Cu^2+^ and Zn^2+^, literature IC_50_ values for dinoflagellate bioluminescence (Lapota et al. 1993, 1995; Heimann et al. 2002; Rosen et al. 2008; Stauber et al. 2008) span ~0.060–3.200 mg L^−1^ for Cu^2+^ and ~0.345–7.000 mg L^−1^ for Zn^2+^, which is above the maximum concentration tested in this study (~0.064 mg L^−1^). Accordingly, higher concentrations would be required here to estimate IC_50_ for these essential metals. Nevertheless, the three highest doses in our experiments produced significant inhibition relative to controls, indicating that *P. lunula* bioluminescence is sensitive to Cu^2^+ and Zn^2^+ even at comparatively low levels.

Published data also indicate species-specific differences: *L. polyedrum* often appears more sensitive to Cu^2+^ than other dinoflagellates, and some algal bioluminescence assays report greater sensitivity to essential metals than to non-essential ones, patterns not observed in our dataset (Trevors et al. 1986; Heimann et al. 2002; Rosen et al. 2008; Stauber et al. 2008; Nowicka 2022; Le et al. 2024).

Bioluminescence-based toxicity is also assessed using the bacterium *A. fischeri* with standardized protocols (15–30 min assays; EPA 2013; ABNT 2021a–c). Comparing bacterial and dinoflagellate IC_50_ values (Fig. 5; Dutka and Kwan 1981; McCloskey et al. 1996; Kungolos et al. 2004; Fulladosa et al. 2005; Petala et al. 2005; Rosen et al. 2008; Teodorović et al. 2009) shows that *P. lunula* (and other dinoflagellates) are markedly more sensitive to Cd^2+^ and Pb^2+^. For *A. fischeri*, Cd^2+^ IC_50_ values range ~10.9–56.8 mg L^−1^ and Pb^2+^ ~0.120–35.970 mg L^−1^, versus ~0.014–1.710 mg L^−1^ (Cd^2+^) and ~0.016– 2.710 mg L^−1^ (Pb^2+^) in dinoflagellates. In relation to essential metals, cross-organism sensitivity is more comparable, though slightly lower for the bacterium: Cu^2+^ ~0.300– 3.800 mg L^−1^ and Zn^2+^ ~0.860–35.000 mg L^−1^ for *A. fischeri*, versus ~0.060–3.200 mg L^−1^ (Cu^2+^) and ~0.345–7.000 mg L^−1^ (Zn^2+^) for bioluminescent algae (Fig. 5).

Dinoflagellate bioluminescence assays also show sensitivity comparable to, or exceeding, that of traditional metazoan models. For *D. magna* (24–48 h assays; Fig. 5), Cd^2+^ sensitivity after 48 h (IC_50_-48 h = 0.170 mg L^−1^; Teodorović et al. 2009) is similar, though slightly lower, than *P. lunula* values at 4 h (0.133 mg L^−1^; Heimann et al. 2002) and 24 h (0.014 mg L^−1^; present study). For Pb^2+^, dinoflagellates are far more sensitive: *D. magna*, IC_50_-24 h = 208.140 mg L^−1^ and IC_50_-48 h = 74.730 mg L^−1^ (Teodorović et al. 2009). For Cu^2+^ and Zn^2+^, sensitivities are broadly similar across organisms, with *D. magna*, IC_50_-24 h of 0.050 mg L^−1^ for Cu^2^+ and ~0.410–1.800 mg L^−1^ for Zn^2+^ (EPA 1980; Kungolos et al. 2004; Teodorović et al. 2009). Acute tests with *H. azteca*, which quantify mortality, indicate even greater sensitivity to all four metals (IC_50_-96 h = 0.001 mg L^−1^ for Cd^2+^, Cu^2+^, and Zn^2+^; 0.005 mg L^−1^ for Pb^2+^; Borgmann et al. 2005; Fig. 5). Even so, dinoflagellate bioluminescence assays detected effects at low doses and offer a rapid, sensitive alternative or complement to traditional ecotoxicological models. Finally, the plate-based bioluminescence approach used here proved highly efficient for detecting Cd^2+^ and Pb^2+^. A modern 96-well luminometer enabled simultaneous analysis of many samples without sacrificing precision, streamlining screening and large-scale evaluations while maintaining data quality.

## CONCLUSION

The results of this study demonstrated a clear dose-dependent effect on the bioluminescence of *P. lunula* in response to the trace metals investigated, highlighting that the proposed bioassay methodology with bioluminescent dinoflagellates and a microplate luminometer significantly improves pollutant detection by employing modern light-measurement equipment. The sensitivity observed in *P. lunula* was comparable to, and in some cases exceeded, that of organisms widely used in ecotoxicology. The 24 h IC_50_ values were 0.014 mg L^−1^ for Cd^2+^ and 0.016 mg L^−1^ for Pb^2+^. In comparison, *A. fischeri* showed IC_50_ values ranging from 10.900–56.800 mg L^−1^ for Cd^2+^ and 0.120– 35.970 mg L^−1^ for Pb^2+^, while *D. magna* exhibited IC_50_ values ranging from 0.170–0.290 mg L^−1^ and 74.730–208.140 mg L^−1^, respectively. These results highlight the strong potential of dinoflagellates in environmental applications, though further testing with essential metals is needed to establish IC_50_ values. Nevertheless, statistical analyses in this study already confirmed dose-dependent inhibition for Cu^2+^ and Zn^2+^, and previous research has likewise shown that dinoflagellates exhibit responses to these metals comparable to those of established ecotoxicological model organisms.

This approach confirms bioluminescent dinoflagellates as practical tools for environmental monitoring, combining low cost, high sensitivity, ease of cultivation, ecological relevance as primary producers, and readily measurable sublethal light responses. These features also make them suitable for preliminary screening. In the face of rising pollution, such methods strengthen monitoring efforts and guide timely management. Their use broadens the ecotoxicology toolkit, supports mitigation before irreversible damage, and aligns with the United Nations Decade of Ocean Science for Sustainable Development.

## Declarations

## Acknowledgments

We thank the Aidar & Kutner Microorganism Bank (Oceanographic Institute, University of São Paulo) for providing dinoflagellate strains, assistance with culture maintenance, and access to facilities.

## Availability of data and materials

Datasets generated and analyzed in this study are available from the corresponding author on request.

## Code availability

Not applicable.

## Authors’ contributions

LSP: data curation, formal analysis, investigation, methodology, writing (original draft, review and editing); FMS: resources, supervision, writing (review and editing); MM: formal analysis, writing (review and editing); LR: formal analysis, writing (review and editing); ESB: investigation, supervision, resources, supervision, writing (review and editing); AGO: conceptualization, funding acquisition, investigation, resources, supervision, writing (review and editing). All authors approved the final manuscript and agreed to its submission.

## REFERENCES

ABNT (2021a) NBR 15411-1: ecotoxicologia aquática – determinação do efeito inibitório de amostras aquosas sobre a emissão de luz de Vibrio fischeri (Ensaio de bactéria luminescente) Parte 1: método utilizando bactérias recém-cultivadas. Rio de Janeiro.

ABNT (2021b) NBR 15411-2: ecotoxicologia aquática – determinação do efeito inibitório de amostras aquosas sobre a emissão de luz de Vibrio fischeri (Ensaio de bactéria luminescente) Parte 2: método utilizando bactérias desidratadas. Rio de Janeiro.

ABNT (2021c) NBR 15411-3: ecotoxicologia aquática – determinação do efeito inibitório de amostras de aquosas sobre a emissão de luz de Vibrio fischeri (Ensaio de bactéria luminescente) Parte 3: método utilizando bactérias liofilizadas. Rio de Janeiro.

Allan R (1997) Introduction: mining and metals in the environment. J Geochem Explor 58:95–100.

Borgmann U, Couillard Y, Doyle P, Dixon DG (2005) Toxicity of sixty-three metals and metalloids to Hyalella azteca at two levels of water hardness. Environ Toxicol Chem 24:641–652.

Claes JM, Haddock SHD, Coubris C, Mallefet J (2024) Systematic Distribution of Bioluminescence in Marine Animals: A Species-Level Inventory. Life 14:432.

Craig JM, Klerks PL, Heimann K, Waits JL (2003) Effects of salinity, pH and temperature on the re-establishment of bioluminescence and copper or SDS toxicity in the marine dinoflagellate Pyrocystis lunula using bioluminescence as an endpoint. Environ Pollut 125:267–275.

Dutka BJ, Kwan KK (1981) Comparison of Three Microbial Toxicity Screening Tests with the Microtox Test. Bull Environ Contam Toxicol 27:753–757.

Fajardo C, Amil-Ruiz F, Fuentes-Almagro C, De Donato M, Martinez-Rodriguez G, Escobar-Niño A, Carrasco R.; Mancera JM, Fernandez-Acero FJ (2019) An “omic” approach to Pyrocystis lunula: New insights related with this bioluminescent dinoflagellate. J Proteomics 209.

Fajardo C, De Donato M, Rodulfo H, Martinez-Rodriguez G, Costas B, Mancera JM, Fernandez-Acero FJ (2020) New perspectives related to the bioluminescent system in dinoflagellates: Pyrocystis lunula, a case study. Int. J Mol Sci 21.

Fávero LP, Belfiore P (2017) Manual da Análise de Dados - Estatística e Modelagem Multivariada com Excel^®^, SPSS^®^ e Stata^®^. Elsevier 1189.

Fulladosa E, Murat JC, Villaescusa I (2005) Study on the toxicity of binary equitoxic mixtures of metals using the luminescent bacteria Vibrio fischeri as a biological target. Chemosphere 58:551–557.

Gerdes HH, Kaether C (1996) Green fluorescent protein applications in cell biology. FEBS Letters 389:44–47.

Girotti S, Ferri EN, Fumo MG, Maiolini E (2008) Monitoring of environmental pollutants by bioluminescent bacteria. Anal Chim Acta 608:2–29.

Green DE, Fry M, Blondin GA (1980) Phospholipids as the molecular instruments of ion and solute transport in biological membranes. Proc Natl Acad Sci USA 77:257–61.

Guillard RLR (1975) Culture of Phytoplankton for Feeding Marine Invertebrates. In: Smith W, Chanley MH (Ed.) Culture of Marine Invertebrates. London, Plenum Press.

Guillard RRL, Ryther JH (1962) Studies of Marine Planktonic Diatoms: Cyclotella nana Hustedt, and Detonula confervaceae (Cleve) Gran. Can J Microbiol 8:229–239.

Haddock SHD, Moline MA, Case JF (2010) Bioluminescence in the Sea. Annu Rev Mar Sci 2:443–493.

Hannan PJ, Stiffey AV, Jarvist BB (1986) Bioluminescence as the Basis for the Detection of Trichothecenes. Naval Research Laboratory Memoradum Report USA.

Hastings JW (2013) Bioluminescence: Living Lights, Lights for Living. United Kingdom, Harvard University.

Heimann K, Matuszewski JM, Klerks PL (2002) Effects of metals and organic contaminants on the recovery of luminescence in the marine dinoflagellate Pyrocystis lunula (Dinophyceae). J Phycol 38:482–492.

Herring PJ (1987) Systematic Distribution of Bioluminescence in Living Organisms. J Biolumin Chemilumin 1:147–163.

Hildenbrand ZL, Osorio A, Carlton DD, Fontenot BE, Walton JL, Hunt LR, Oka H, Hopkins D, Bjorndal B, Schug KA (2015) Rapid analysis of eukaryotic bioluminescence to assess potential groundwater contamination events. J Chem.

Islam MS, Tanaka M (2004) Impacts of pollution on coastal and marine ecosystems including coastal and marine fisheries and approach for management: a review and synthesis. Mar Pollut Bull 48:624–649.

Kungolos A, Hadjispyrou S, Petala M, Tsiridis V, Samaras P, Sakellaropoulos GP (2004) Toxic properties of metals and organotin compounds and their interactions on Daphnia magna and Vibrio fischeri. Water Air Soil Poll. Focus 4:101–110.

Lapota D, Duckworth D, Rosenberger DE, Copeland HD, Mastny GF (1995) The QWIKLITE Bioluminescence Bioassay System to Assess Toxic Effects in the Biosphere. Environmental Quality Technology: Advancing the Pillars Toward the21st Century pp 153–167.

Lapota D, Moskowitz GJ, Rosenberger DE, Grovhoug JG (1993) The use of stimulable bioluminescence from marine dinoflagellates as a means of detecting toxicity in the marine environment. Environmental Toxicology and Risk Assessment.

Lau ES, Oakley TH (2021) Multi-level convergence of complex traits and the evolution of bioluminescence. Biol Rev 96:673–691.

Le VG, Nguyen MK, Nguyen HL, Thai VA, L. VR, Vu QM, Asaithambi P, Chang SW, Nguyen DD (2024) Ecotoxicological response of algae to contaminants in aquatic environments: a review. Environ Chem Lett 22:919–939.

Magalhães DP, Ferrão-Filho AS (2008) A ecotoxicologia como ferramenta no biomonitoramento de ecossistemas aquáticos. Oecol Bras 3:355–381.

Mccloskey JT, Newman MC, Clark SB (1996) Predicting the relative toxicity of metal ions using ion characteristics: Microtox® bioluminescence assay. Environ Toxicol Chem 15:1730–1737.

Moline MA, Oliver MJ, Mobley CD, Sundman L, Bensky T, Bergmann T, Bissett WP, Case J, Raymond EH, Schofield OME (2007) Bioluminescence in a complex coastal environment: 1. Temporal dynamics of nighttime water-leaving radiance. J. Geophys. Res. Oceans 112.

Nowicka B (2022) Heavy metal–induced stress in eukaryotic algae—mechanisms of heavy metal toxicity and tolerance with particular emphasis on oxidative stress in exposed cells and the role of antioxidant response. Environ Sci Pollut Res 29:16860– 16911.

Ogawa H, Inouye S, Tsuji FI, Yasuda K, Umesono K (1995) Localization, trafficking, and temperature-dependence of the Aequorea green fluorescent protein in cultured vertebrate cells. Proc Natl Acad Sci USA 92:11899–11903.

Okamoto OK, Shao L, Hastings JW, Colepicolo P (1999) Acute and chronic effects of toxic metals on viability, encystment and bioluminescence in the dinoflagellate Gonyaulax polyedra. Comp. Biochem. Physiol. C 123:75–83.

Omar HH (2002) Bioremoval of zinc ions by Scenedesmus obliquus and Scenedesmus quadricauda and its effect on growth and metabolism. Int Biodeterior Biodegrad 50:95–100.

Perin LS, Moraes GV, Galeazzo GA, Oliveira AG (2022) Bioluminescent Dinoflagellates as Bioassay for Toxicity Assessment. Int J Mol Sci 23:13012.

Perkink EJ (1979) The need for sublethal studies. Phil Trans R Soc Lond B 286: 425–442.

Petala M, Tsiridis V, Kyriazis S, Samaras P, Kungolos A, Sakellaropoulos G P (2005) Evaluation of toxic response of heavy metals and organic pollutants using Microtox acute toxicity test. 9th International Conference on Environmental Science and Technology, Rhodes Island, Greece.

Pinto E, Sigaud-Kutner TCS, Leitão MAS, Okamoto OK, Morse D, Colepicolo P (2003) Heavy metal–induced oxidative stress in algae. J Phycol 39:1008–1018.

Rand GM (1995) Fundamentals of aquatic toxicology: effects, environmental fate and risk assessment. United States, CRC press.

Raven JA, Evans MCW, Korb RE (1999) The role of trace metals in photosynthetic electron transport in O_2_-evolving organisms. Photosynth Res 60:111–150.

Rosen G, Osorio-Robayo A, Rivera-Duarte I, Lapota D (2008) Comparison of bioluminescent dinoflagellate (Qwiklite) and bacterial (Microtox) rapid bioassays for the detection of metal and ammonia toxicity. Arch Environ Contam Toxicol 54:606–611.

Schramm S, Weiß D (2024) Bioluminescence – The Vibrant Glow of Nature and its Chemical Mechanisms. ChemBioChem 25.

Schoofs H, Schmit J, Rink L (2024) Zinc Toxicity: Understanding the Limits. Molecules 29:3130.

Sebaugh JL (2011) Guidelines for accurate EC50/IC50 estimation. Pharm Stat 10:128–134.

Shimomura O (2006) Bioluminescence - Chemical Principles and Methods. Singapore, World Scientific Publishing Company.

Sharma RK, Agrawal M (2005) Biological effects of heavy metals: An overview. J Environ Biol 26:301–313.

Singh S, Parihar P, Singh R, Singh VP, Prasad SM (2016) Heavy Metal Tolerance in Plants: Role of Transcriptomics, Proteomics, Metabolomics, and Ionomics. Front Plant Sci 6:1143.

Stauber JL, Binet MT, Bao VWV, Boge J, Qzhang A, Leung MEUK, Adams MS (2008) Comparison of the QwikLiteTM algal bioluminescence test with marine algal growth rate inhibition bioassays. Environ Toxicol 23:617–625.

Stern BR (2010) Essentiality and Toxicity in Copper Health Risk Assessment: Overview, Update and Regulatory Considerations. J Toxicol Environ Health A 73:114–127.

Syed AJ, Anderson JC (2021) Applications of bioluminescence in biotechnology and beyond. Chem Soc Rev 50:5668–5705.

Taylor FJR (1987) The Biology of Dinoflagellates. United States, Wiley–Blackwell.

Teodorovic I, Planojevic I, Knezevic P, Radak S, Nemet I (2009) Sensitivity of bacterial vs. acute Daphnia magna toxicity tests to metals. Cent Eur J Biol 4:482–492.

Timsit Y, Lescot M, Valiadi M, Not F (2021) Bioluminescence and Photoreception in Unicellular Organisms: Light-Signalling in a Bio-Communication Perspective. Int J Mol Sci 22:11311.

Travieso L, Pellón A, Benítez F, Sánchez E, Borja R, O’farrill N, Weiland P (2002) Bioalga reactor: preliminary studies for heavy metals removal. Biochem Eng J 12:87– 91.

Trevors JT, Stratton GW, Gadd GM (1986) Cadmium transport, resistance, and toxicity in bacteria, algae, and fungi. Can J Microbiol 32:447–64.

Truhaut R (1977) Ecotoxicology: objectives, principles and perspectives. Ecotoxicol Environ Saf 1:151–173.

U.S. Environmental Protection Agency (1980) Ambient waterquality criteria for zinc. Washington, DC: U.S. EPA.

U.S. Environmental Protection Agency (2013) Strategic Diagnostics Inc. Microtox® Rapid Toxicity Testing System. Environmental Technology Verification. Washington, DC: U.S. EPA.

Vinutha HP, Poornima B, Sagar BM (2018) Detection of Outliers Using Interquartile Range Technique from Intrusion Dataset. Singapore, Springer Nature.

Valiadi M, Iglesias-Rodriguez D (2013) Understanding bioluminescence in dinoflagellates—how far have we come? Microorganisms 1: 3–25.

Visbeck M (2018) Ocean science research is key for a sustainable future. Nat Commun 9:690.

Wilson T, Hastings JW (1998) Bioluminescence. Annu Rev Cell Dev Biol 14:197–230.

Zagatto A, Bertoletti E (2006) Ecotoxicologia aquática-princípios e aplicações. Brazil, RiMa.

