## Supplementary Information for "High-throughput microplate luminometry of *Pyrocystis lunula* bioluminescence for metal toxicity assessment"

| **a** | 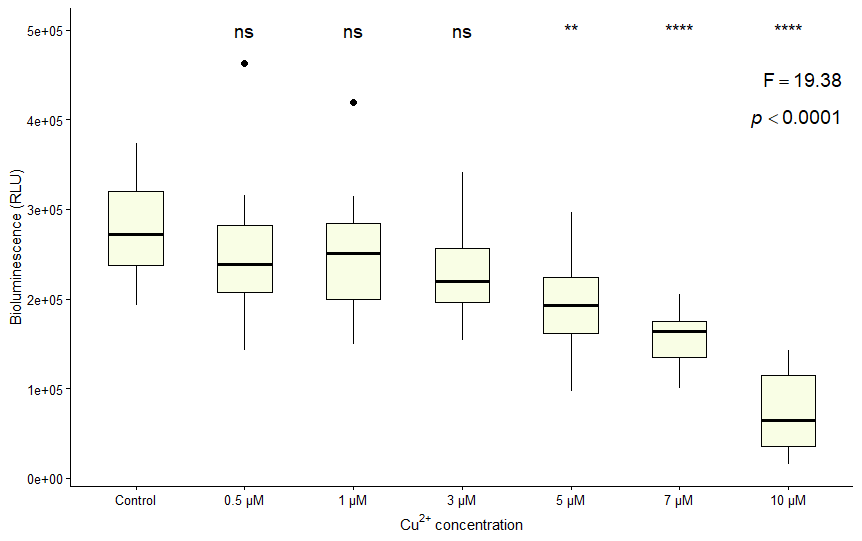 |
| --- | --- |
| **b** | 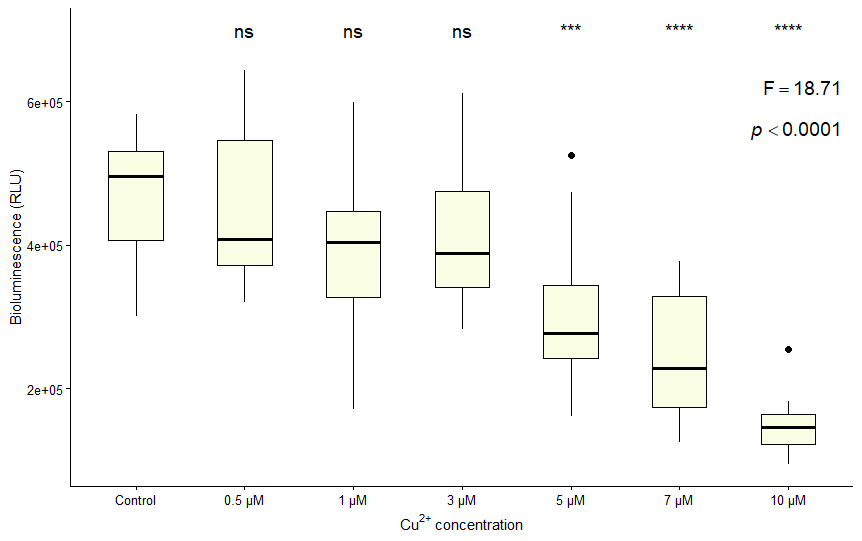 |
| **c** | 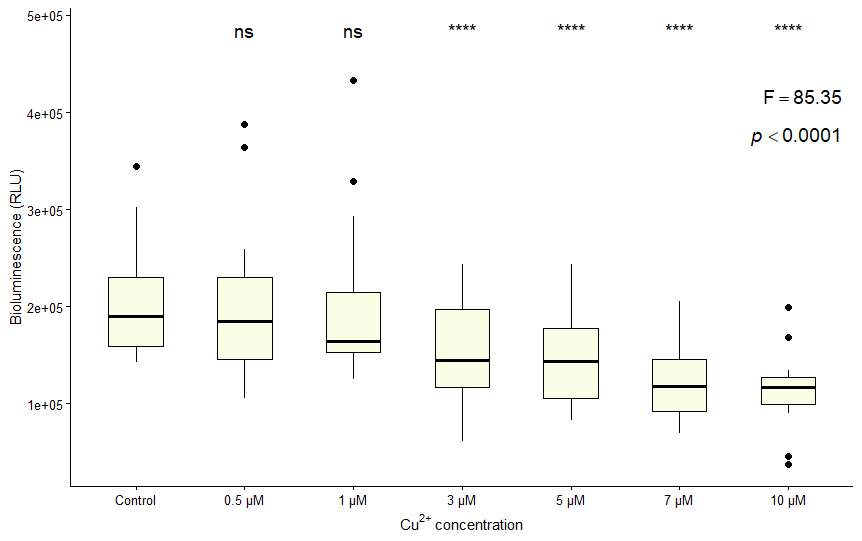 |

**Fig. S1** Light emission of *P. lunula* at different concentrations of Cu^2+^. **a**) replicate 1; **b**) replicate 2; **c**) replicate 3. (****) indicates a significant difference compared to the control (**p **< 0.02); (***)** indicates a significant difference compared to the control (p ≤ 0.0005); (****) indicates a significant difference compared to the control (p < 0.0001).

| **a** | 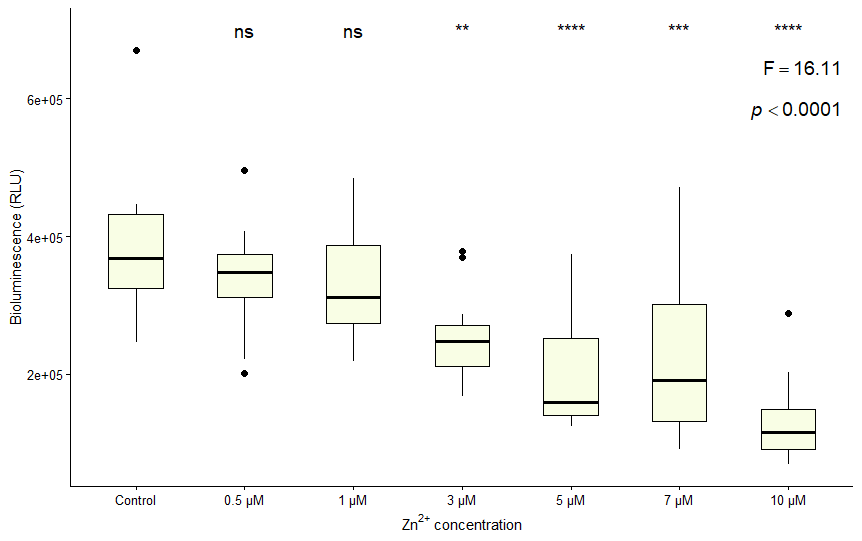 |
| --- | --- |
| **b** | 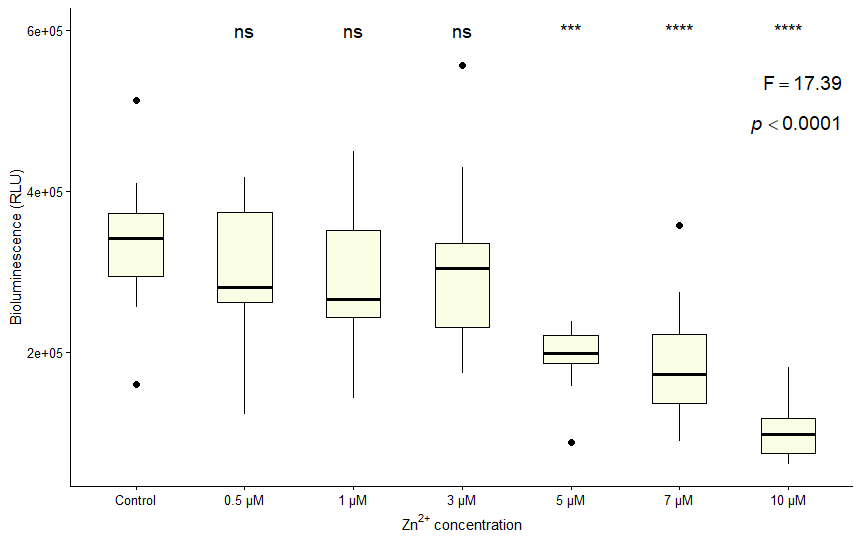 |
| **c** | 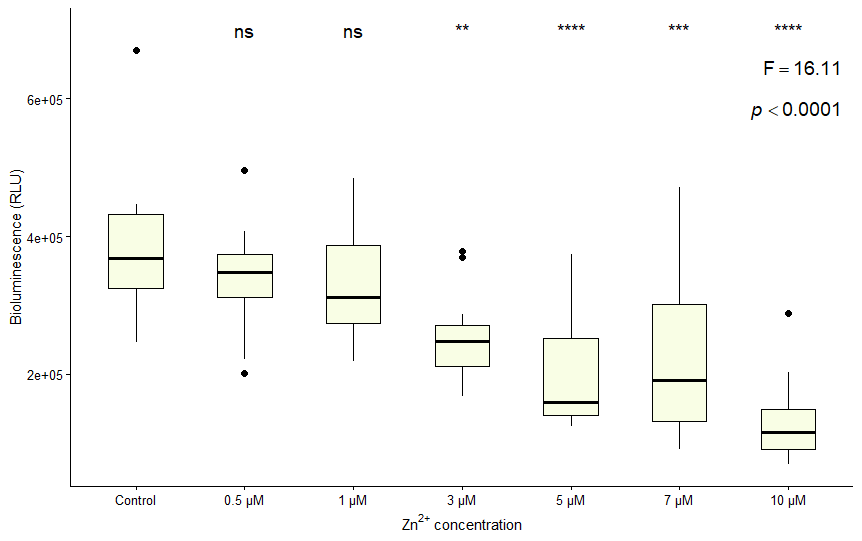 |

**Fig. S2** Light emission of *P. lunula* at different concentrations of Zn^2+^. **a**) replicate 1; **b**) replicate 2; **c**) replicate 3. (****) indicates a significant difference compared to the control (**p **< 0.03); (***)** indicates a significant difference compared to the control (p ≤ 0.0005); (****) indicates a significant difference compared to the control (p < 0.0001).

| **a** | 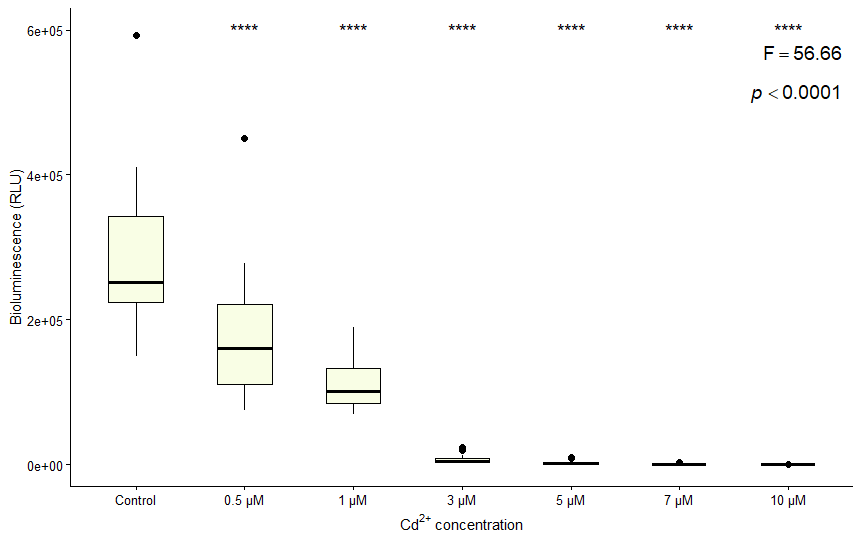 |
| --- | --- |
| **b** | 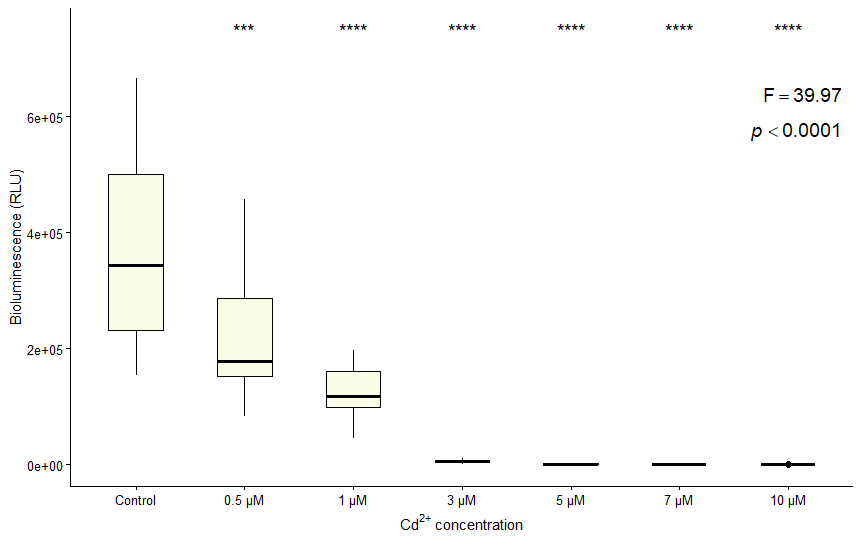 |
| **c** | '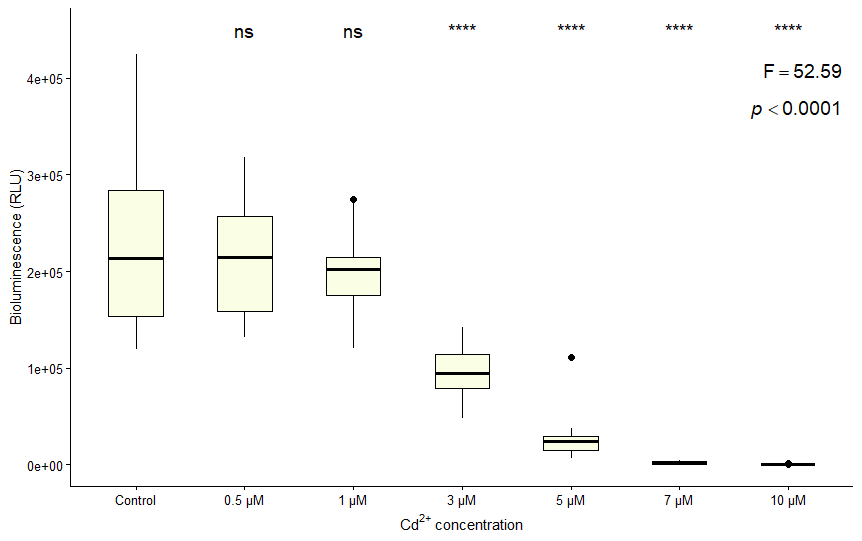 |

**Fig. S3** Light emission of *P. lunula* at different concentrations of Cd^2+^. **a**) replicate 1; **b**) replicate 2; **c**) replicate 3. **(***)** indicates a significant difference compared to the control (p ≤ 0.0003); (****) indicates a significant difference compared to the control (p < 0.0001).

| **a** | 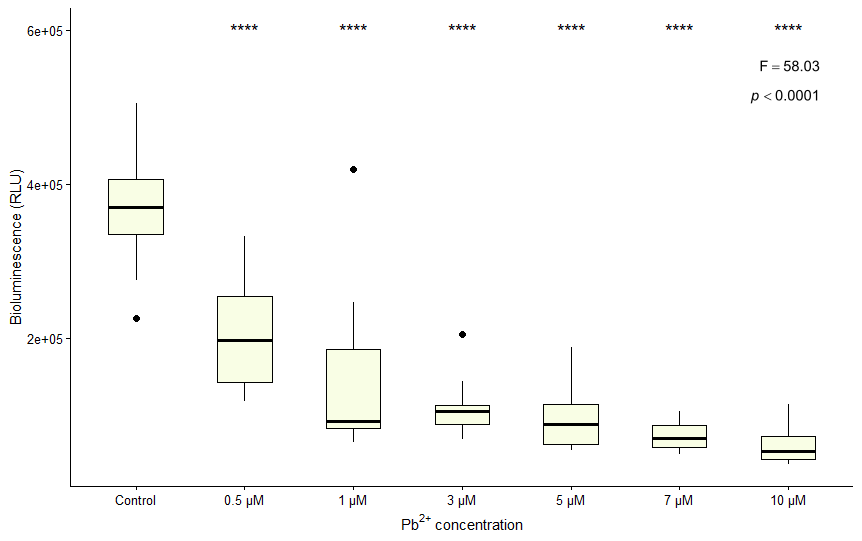 |
| --- | --- |
| **b** | 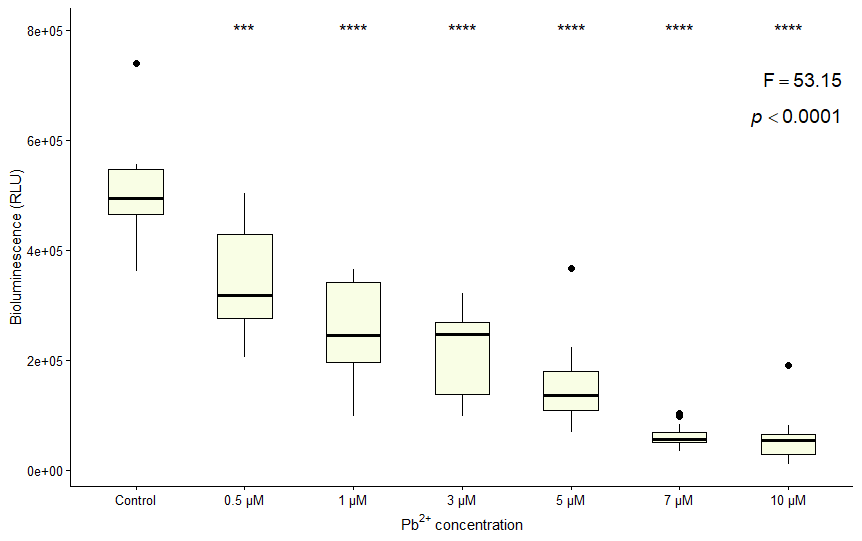 |
| **c** | 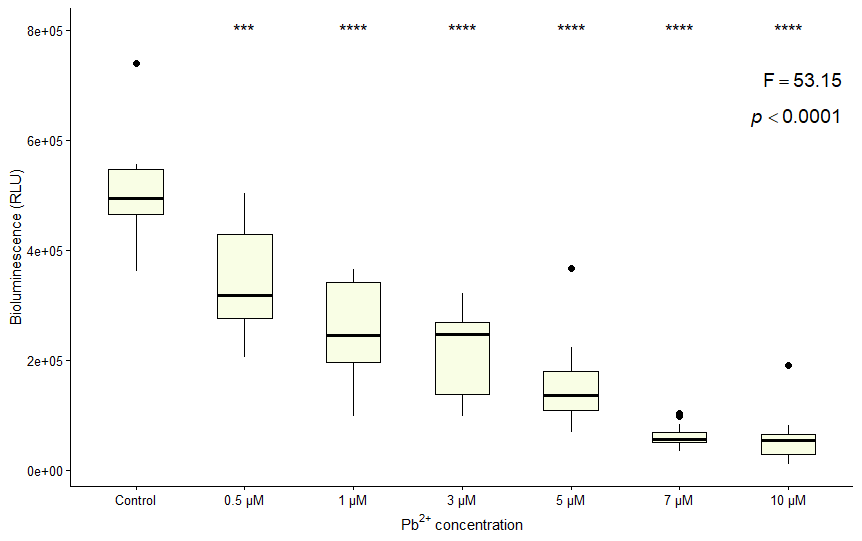 |

**Fig. S4** Light emission of *P. lunula* at different concentrations of Pb^2+^. **a**) replicate 1; **b**) replicate 2; **c**) replicate 3. **(***)** indicates a significant difference compared to the control (p ≤ 0.001); (****) indicates a significant difference compared to the control (p < 0.0001).
